# Binette: a fast and accurate bin refinement tool to construct high quality Metagenome Assembled Genomes

**DOI:** 10.1101/2024.04.20.585171

**Authors:** Jean Mainguy, Claire Hoede

## Abstract

Metagenomics enables the study of microbial communities and their individual members through shotgun sequencing. An essential phase of metagenomic analysis is the recovery of metagenome-assembled genomes (MAGs). In a metagenomic analysis, sequence reads are assembled into contigs, which are then grouped into bins based on common characteristics - a process known as binning - to generate MAGs. The approach of applying multiple binning methods and combining them in a process called bin refinement allows us to obtain more and higher quality MAGs from metagenomic datasets. We present Binette, a bin refinement tool inspired by metaWRAP’s bin refinement module, which addresses the limitations of the latter and ensures better results. Binette achieves this by creating new hybrid bins using basic set operations from the input bin sets. CheckM2 is then used to assess bin quality and select the best possible bins.

## Statement of need

Metagenomics enables the study of microbial communities and their individual members through shotgun sequencing. An essential phase of metagenomic analysis is the recovery of metagenome-assembled genomes (MAGs). MAGs serve as a gateway to additional analyses, including the exploration of organism-specific metabolic pathways, and form the basis for comprehensive large-scale metagenomic surveys [Acinas et al., 2021, Nayfach et al., 2019].

In a metagenomic analysis, sequence reads are first assembled into longer sequences called contigs. These contigs are then grouped into bins based on common characteristics in a process called binning to obtain MAGs. There are several tools that can be used to bin contigs into MAGs. These tools are based on various statistical and machine learning methods and use contig characteristics such as tetranucleotide frequencies, GC content and similar abundances across samples [Alneberg et al., 2014, Kang et al., 2019, Nissen et al., 2021].

The approach of applying multiple binning methods and combining them has proven useful to obtain more and better quality MAGs from metagenomic datasets.This combination process is called bin-refinement and several tools exist to perform such tasks, such as DAS Tool [Sieber et al., 2018], MagScot [Rühlemann et al., 2022] and the bin-refinement module of the metaWRAP pipeline [Uritskiy et al., 2018]. Of these, metaWRAP’s bin-refinement tool has demonstrated remarkable efficiency in benchmark analysis [Meyer et al., 2022].

However, it has certain limitations, most notably its inability to integrate more than three binning results. In addition, it repeatedly uses CheckM [Parks et al., 2015] to assess bin quality throughout its execution, which contributes to its slower performance. Furthermore, since it is embedded in a larger framework, it may present challenges when attempting to integrate it into an independent analysis pipeline.

We present Binette, a bin refinement tool inspired by metaWRAP’s bin refinement module, which addresses the limitations of the latter and ensures better results.

## Summary

Binette is a Python reimplementation and enhanced version of the bin refinement module used in metaWRAP. It takes as input sets of bins generated by various binning tools. Using these input bin sets, Binette constructs new hybrid bins using basic set operations. Specifically, a bin can be defined as a set of contigs, and when two or more bins share at least one contig, Binette generates new bins based on their intersection, difference, and union (Figure 1.A). This approach differs from metaWRAP, which exclusively generates hybrid bins based on bin intersections and allows Binette to expand the range of possible bins.

**Figure 1:**
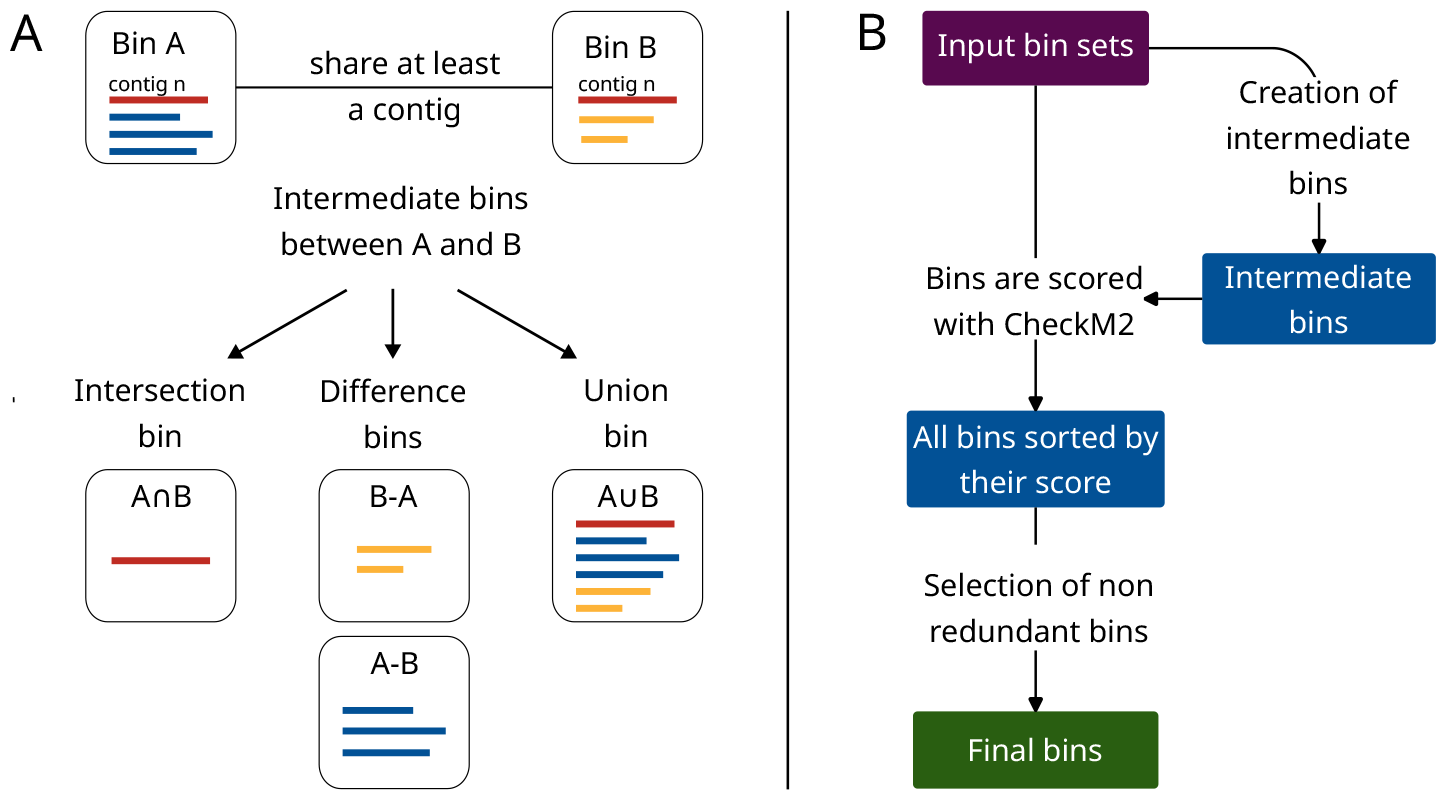
Overview of Binette Steps. (A) Intermediate Bin Creation Example: Bins are represented as square shapes, each containing colored lines representing the contigs they contain. Creation of intermediate bins involves the initial bins sharing at least one contig. Set operations are applied to the contigs within the bins to generate these intermediate bins. **(B) Binette Workflow Overview**: Input bins serve as the basis for generating intermediate bins. Each bin undergoes a scoring process utilizing quality metrics provided by CheckM2. Subsequently, the bins are sorted based on their scores, and a selection process is executed to retain non-redundant bins.

Bin completeness and contamination are assessed using CheckM2 [Chklovski et al., 2023]. Bins are scored using the following scoring function: *completeness* − *weight* * *contamination*, with the default weight set to 2. These scored bins are then sorted, facilitating the selection of a final new set of non-redundant bins (Figure 1.B). The ability to score bins is based on CheckM2 rather than CheckM1 as in the metaWRAP pipeline. CheckM2 uses a novel approach to evaluate bin quality based on machine learning techniques. This approach improves speed and also provides better results than CheckM1. Binette initiates CheckM2 processing by running its initial steps once for all contigs within the input bins. These initial steps involve gene prediction using Prodigal and alignment against the CheckM2 database using Diamond [Buchfink et al., 2015]. Binette uses Pyrodigal [Larralde, 2022], a Python module that uses Cython to provide bindings to Prodigal [Hyatt et al., 2010]. The intermediate Checkm2 results are then used to assess the quality of individual bins, eliminating redundant calculations and speeding up the refinement process.

Binette serves as the bin refinement tool within the metagWGS metagenomic analysis pipeline [Mainguy et al., prep], providing a robust and faster alternative to the bin refinement module of the metaWRAP pipeline as well as other similar bin refinement tools.

## Benchmark

We have compared Binette with DAS Tool [Sieber et al., 2018] using the simulated mouse gut metagenome data released in preparation for the second round of CAMI II challenges [Meyer et al., 2022]. Both tools were compared by following the binning benchmark described in this Nature Protocols article [Meyer et al., 2021].

As input of the bin refinement tools, we used the results produced from the cross-sample gold-standard assembly with the binning tools MaxBin v.2.2.7 [Wu et al., 2016], MetaBAT v.2.12.1 [Kang et al., 2019], and CONCOCT v.1.0.0 [Alneberg et al., 2014] available in the CAMI tool result repositories on Zenodo. Binning results from the individual binners and refinement tools have been evaluated using AMBER v2.0.4 [Meyer et al., 2018], which generated binning quality metrics based on the simulated data’s ground truth.

Although Concoct and MaxBin have the highest average completeness, this advantage is offset by very low average purity. Metabat2, on the other hand, has a low average completeness. In contrast, Binette and DAS Tool show comparable and superior results in both average completeness and purity, outperforming individual binners (Suppl. Table 1).

In the context of recovering high quality genomes (completeness > 90% and contamination < 5%), Binette outperformed DAS Tool, recovering 367 genomes compared to DAS Tool’s 334 high quality genomes. Notably, individual binning tools had a lower ability to recover high-quality genomes compared to the refinement tools (Figure 2).

**Figure 2:**
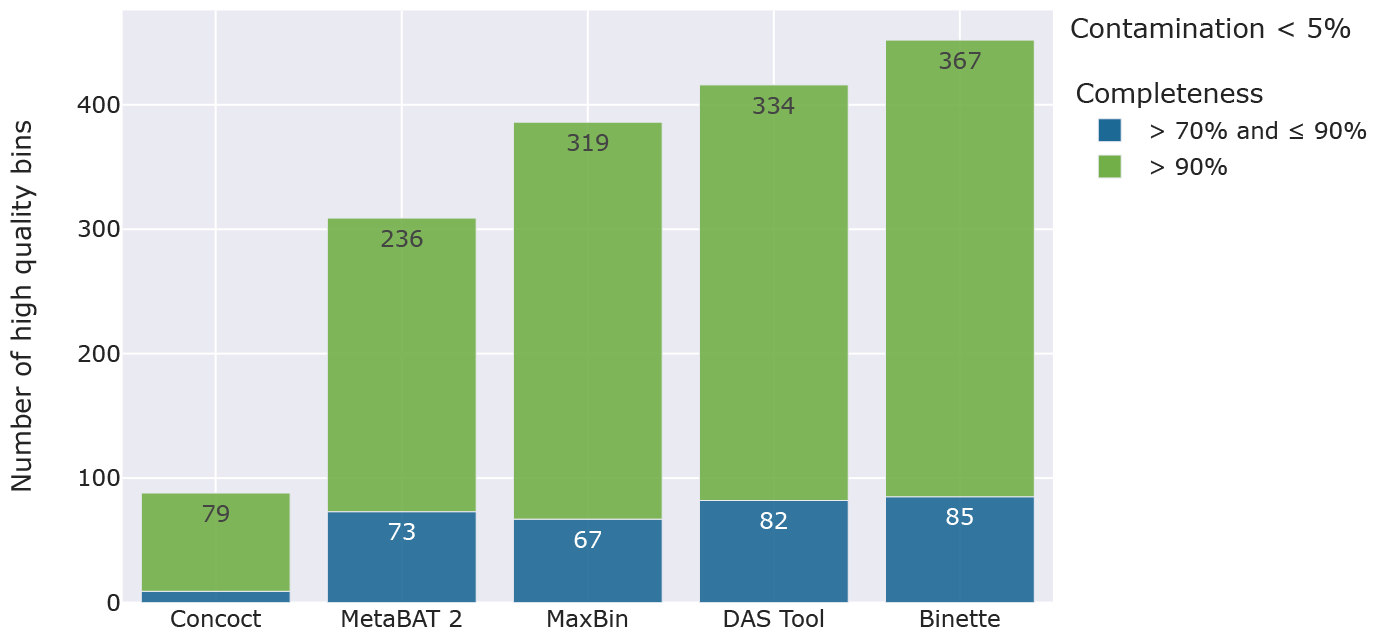
Evaluation of different binning strategies on simulated mouse gut metagenome data from the CAMI II challenges. The bar plot displays the count of high-quality bins with contamination < 5% and completeness > 70% and > 90%. DAS Tool and Binette results were generated based on bins from Concoct, MetaBAT 2, and MaxBin.

These results highlight the importance of using refinement tools to improve overall binning results. While DAS Tool and Binette show comparable performances based on average bin statistics, Binette outperforms DAS Tool in recovering high quality genomes.

## Availability

Binette is available on PyPI and can be installed using standard Python package management tools. Additionally, a dedicated Conda package is available in the Bioconda channel [Grüning et al., 2018]. The source code for Binette is available on GitHub under the MIT license. The GitHub repository includes continuous integration tests, test coverage, and employs continuous deployment through GitHub actions to maintain a robust and reliable codebase.

## Acknowledgements

We would like to thank Matthias Zytnicki for his valuable insights and support during the development of the binette algorithm.

