## Supplemental Table 1 for "Binette: a fast and accurate bin refinement tool to construct high quality Metagenome Assembled Genomes"

| Tool | Average purity (bp) | Average completeness (bp) | Purity (bp) | Completeness (bp) | Adjusted Rand index (seq) | Adjusted Rand index (bp) | Percentage of binned bp | Accuracy |
|---|---|---|---|---|---|---|---|---|
| Gold standard | 1 | 1 | 1 | 1 | 1 | 1 | 1 | 1 |
| Binette 1.0.0 | 0.9265 | 0.6863 | 0.9281 | 0.7076 | **0.8236** | **0.9197** | 0.7463 | 0.6927 |
| DasTool 1.1.2 | **0.9297** | 0.6543 | **0.9317** | 0.6714 | 0.7892 | 0.9137 | 0.7044 | 0.6562 |
| Concoct 1.0.0 | 0.5946 | **0.8469** | 0.4523 | **0.8843** | 0.4320 | 0.4208 | **0.9383** | 0.4244 |
| MaxBin 2.2.7 | 0.7743 | 0.7033 | 0.8172 | 0.7290 | 0.4630 | 0.7578 | 0.8975 | **0.7334** |
| MetaBAT 2.12.1 | 0.9092 | 0.5613 | 0.8500 | 0.5918 | 0.6426 | 0.7252 | 0.6299 | 0.536 |

**Suppl. Table 1** : Quality metrics of different binning tools from the gold-standard assembly of the CAMI II mouse gut dataset. Concoct, MaxBin, and MetaBat2 are individual binning tools and their bins have been refined by DAS Tool and Binette. Metrics presented in this table have been produced by AMBER v2.0.4. The best result of each metric is indicated in bold.
